# Multi-target genome editing reduces polyphenol oxidase activity in wheat (*Triticum aestivum* L.) grains

**DOI:** 10.1101/2023.05.30.542859

**Authors:** Forrest Wold-McGimsey, Caitlynd Krosch, Rocío Alarcón Reverte, Karl Ravet, Andrew Katz, John Stromberger, Richard Esten Mason, Stephen Pearce

## Abstract

Polyphenol oxidases (PPO) are dual activity metalloenzymes that catalyse the production of quinones. In plants, PPO activity may contribute to biotic stress resistance and secondary metabolism but is undesirable for food producers because it causes the discolouration and changes in flavour profiles of products during post-harvest processing. In wheat (*Triticum aestivum* L.), PPO released from the aleurone layer of the grain during milling results in the discolouration of flour, dough, and end-use products, reducing their value.

Loss-of-function mutations in the *PPO1* and *PPO2* paralogous genes on homoeologous group 2 chromosomes confer reduced PPO activity in the wheat grain but limited natural variation and small intergenic distances between these genes complicates the selection of extremely low-PPO wheat varieties by recombination.

In the current study, a CRISPR/Cas9 construct with one single guide RNA (sgRNA) targeting a conserved copper binding domain was used to edit all seven *PPO1* and *PPO2* genes in the spring wheat cultivar ‘Fielder’. Five of the seven edited T_1_ lines exhibited significant reductions in PPO activity, and T_2_ lines had PPO activity up to 86.7% lower than wild-type controls. In the elite winter wheat cultivars ‘Guardian’ and ‘Steamboat’, which have five *PPO1* and *PPO2* genes, PPO activity was reduced by >90% in both T_1_ and T_2_ lines. This study demonstrates that multi-target editing at late stages of variety development could complement selection for beneficial alleles in crop breeding programmes by inducing novel variation in loci inaccessible to recombination.

## INTRODUCTION

Polyphenol oxidases (PPO) are di-copper metalloenzymes found in all land plants except the Arabidopsis genus (Tran et al., 2012). PPOs are dual activity enzymes, catalysing the hydroxylation of monophenols to diphenols (tyrosinase activity, Enzyme Commission (EC) 1.14.18.1) and the oxidation of *o*-diphenols to *o*-quinones (catechol oxidase activity, EC 1.10.3.1) (van Gelder et al., 1997). Quinones react non-enzymatically with cellular thiol and amine groups to produce melanin pigments, causing browning and discolouration of plant tissues. The active site in the PPO proteins for these reactions includes two highly conserved copper binding domains (CuA and CuB) each with three histidine residues that coordinate interactions between phenols and molecular oxygen (Demeke and Morris, 2002). While their physiological function remains unclear, there is indirect evidence that PPO contributes to biotic stress resistance. Many PPO proteins are localized in the chloroplast and come into contact with their phenolic substrates only following senescence, wounding, or physical disruption. In several plant species, *PPO* genes are upregulated in response to wounding or pathogen infection, and variation in PPO activity is associated with resistance to bacterial and fungal pathogens (Zhang and Sun, 2021). PPO may also play a role in plant secondary metabolism (Araji et al., 2014; Sullivan, 2014).

For the food industry, PPO activity is generally undesirable because it causes the discolouration of plant tissues and changes in flavour profile during post-harvest processing. A readily observed example is the browning of fresh fruit and vegetables following cutting. In common wheat (*Triticum aestivum* L.) PPO enzymes released from the aleurone layer of the grain during milling catalyse biochemical reactions that result in the time-dependent darkening and discolouration of flour, dough, and end-use products such as noodles, an undesirable trait for consumers (Taranto et al., 2017). Although this can be mitigated by reducing the flour extraction rate during milling or by using food additives, a more cost-effective approach is to breed wheat varieties with low PPO activity in their grains.

PPO activity in the grain is an amenable trait for wheat breeders, with a broad sense heritability of 0.97 (Baik et al., 1994; Liu et al., 2020). Genetic linkage and association studies consistently find that homoeologous loci on group 2 chromosomes are the most important sources of genetic variation for PPO grain activity (Beecher et al., 2012; Liu et al., 2020; Zhai et al., 2020). Underlying these loci are paralogous *PPO1* and *PPO2* genes that encode PPO enzymes expressed during grain development (Liu et al., 2020). The genome of the wheat landrace ‘Chinese Spring’ contains two *PPO1* paralogs on chromosome 2B (*PPO1-B1* and *PPO1-B2*), giving seven *PPO1* and *PPO2* genes in total at these loci. On each chromosome, these genes are separated by short physical distances (Figure 1). The number of *PPO1* and *PPO2* genes on chromosome 2B ranges from one to three in different wheat varieties (Table S1).

**Figure 1:**
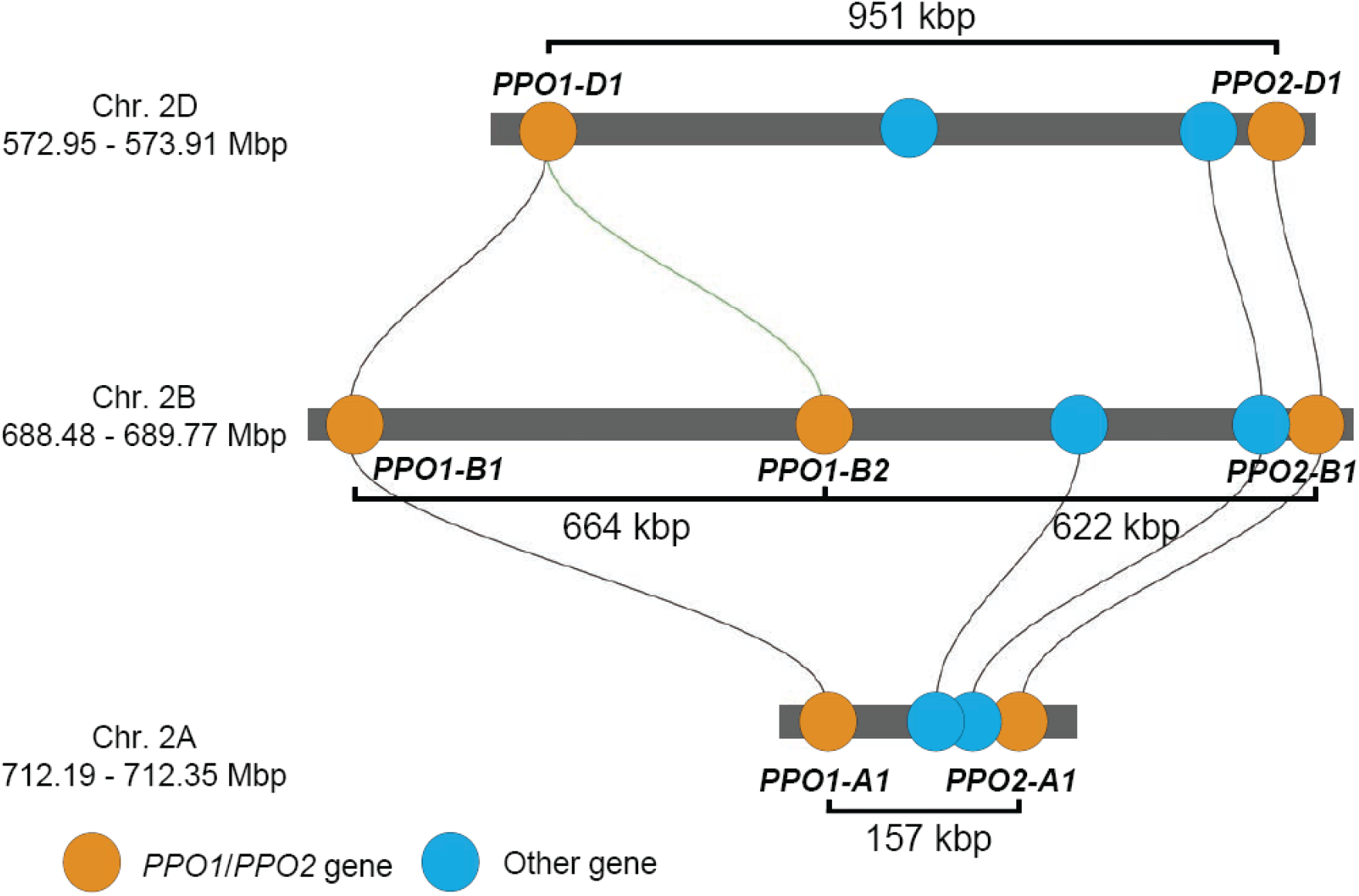
Position of *PPO1* and *PPO2* genes on homoeologous group 2 chromosomes of the wheat landrace ‘Chinese Spring’ IWGSC RefSeq v1.1 genome assembly. Gene positions are drawn to scale and homologous genes linked by lines determined using the Triticeae Gene Tribe microhomology tool (Chen et al., 2020). Physical distances between the start of each gene are labelled. *PPO* genes are named according to guidelines endorsed by the Wheat Initiative (Boden et al., 2023).

Breeding programs can use marker assisted selection to introgress null alleles such as *ppo-A1i* and *ppo-D1c* that confer reduced PPO activity in the wheat grain (He et al., 2007; Hystad et al., 2015). However, to date no natural null alleles have been described for *PPO2-A1*, *PPO2-D1* or for any of the *PPO1* or *PPO2* genes on chromosome 2B that also contribute to PPO activity (Beecher et al., 2012; Liu et al., 2020; Taranto, 2015). In addition to the close physical distances between genes at these loci, this limited natural variation complicates the recombination of non-functional natural variants for each *PPO1* and *PPO2* gene using marker assisted selection.

The genome editing tool CRISPR/Cas9 is now used routinely to induce novel variation at specific genetic loci in crop genomes (Gao, 2021). This technology is particularly useful when multiple simultaneous gene knockouts are required. In the current study, a CRISPR/Cas9 construct with one sgRNA was used to edit all seven *PPO1* and *PPO2* genes in the spring wheat variety ‘Fielder’, conferring significant reductions in PPO grain activity in the T_1_ and T_2_ generation. In two elite winter wheat varieties, PPO activity was reduced by more than 90%. This study demonstrates that carefully designed CRISPR/Cas9 constructs can be used to edit multi-gene families in polyploid crop species and that direct editing of beneficial alleles during the late stages of elite variety development could complement traditional breeding methods for crop improvement.

## RESULTS

### *PPO1* and *PPO2* genes are highly expressed in developing wheat grains

An analysis of a developmental RNA-seq dataset from the wheat landrace ‘Chinese Spring’ showed that among the 20 *PPO* genes, the seven paralogous *PPO1* and *PPO2* genes on group 2 chromosomes are predominantly expressed in developing grain tissues (Figure 2). Some *PPO1* and *PPO2* genes were also expressed in other plant tissues; *PPO1-D1* was highly expressed in stem and spike tissues during anthesis while *PPO1-B1* transcripts were detected in leaf tissues post-anthesis (Figure 2). By contrast, transcript levels of other members of the *PPO* family were low in the developing grain and were more highly expressed in vegetative tissues (Figure 2). These results are consistent with previous studies (Liu et al., 2020), demonstrating that wheat *PPO* genes are developmentally regulated and that *PPO1* and *PPO2* genes contribute the majority of *PPO* transcripts in the grain.

**Figure 2:**
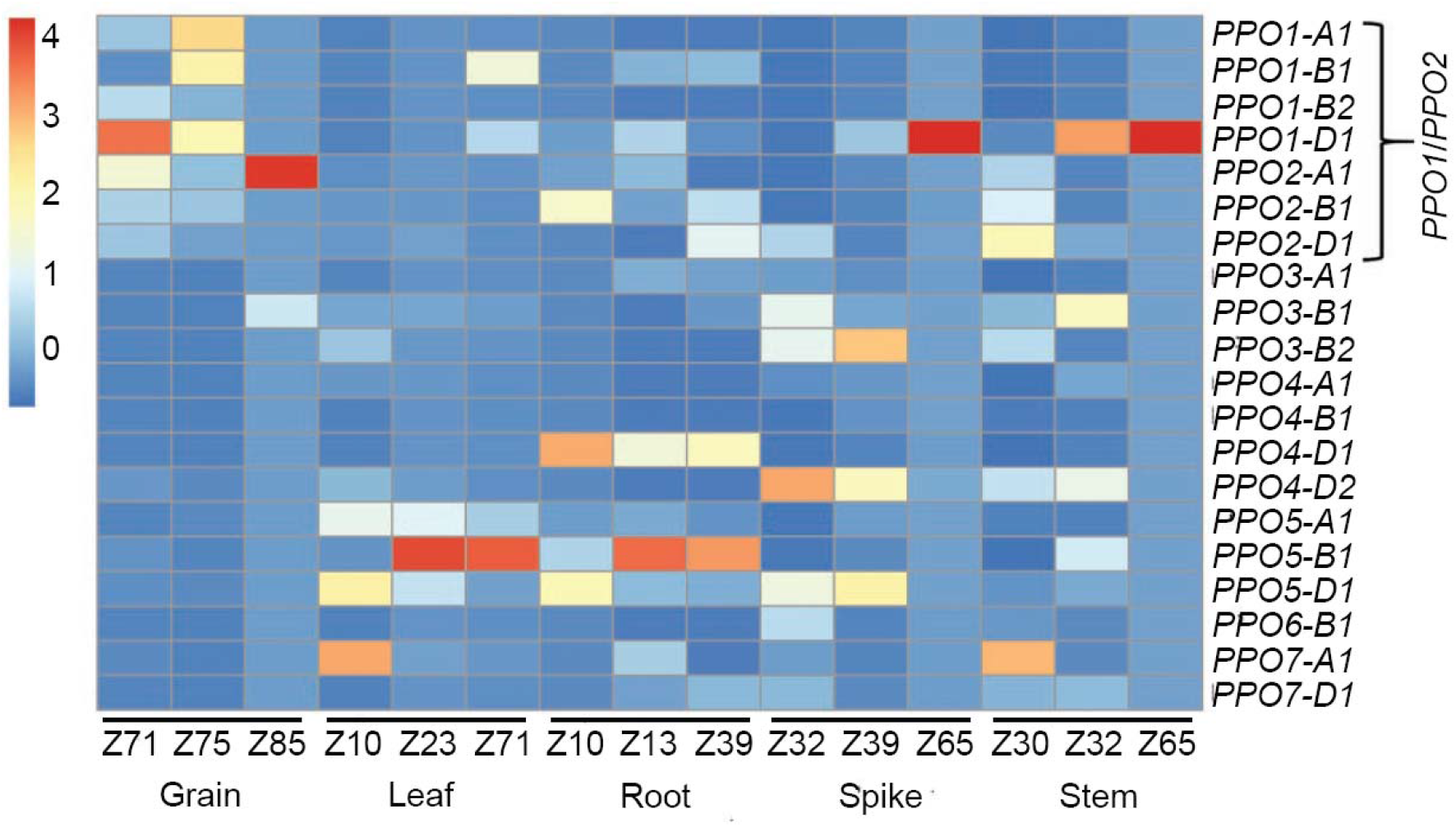
Expression profiles of 20 wheat PPO genes in different wheat tissues. RNA-seq reads from a hexaploid wheat developmental timecourse (Choulet, et al., 2014) were mapped to the IWGSC RefSeq v1.1 wheat reference genome. The developmental stage in each tissue is presented in the Zadoks scale (Zadoks et al., 1974). Expression in transcript per million (TPM) values are scaled for each timepoint using the scale(Data_num) function in the R package pheatmap.

### Multi-target *PPO1* and *PPO2* editing confers significant reductions in PPO grain activity

Based on their expression profile and known association with PPO activity in the grain, all seven *PPO1* and *PPO2* genes were targeted for knockout by genome editing in the spring wheat variety ‘Fielder’. The ‘Fielder’ genome contains 25 *PPO* genes (defined by the presence of tyrosinase, DWL and KWDV domains in their encoded proteins), including all 20 *PPO* genes described in ‘Chinese Spring’ and expansions in *PPO* gene number on chromosomes 3A and 6B (Table S2). Note that *PPO* genes have been named based on their phylogenetic relationships in accordance with guidelines endorsed by the Wheat Initiative (Boden et al., 2023) and do not necessarily match earlier publications. Of the seven *PPO1* and *PPO2* genes in ‘Fielder’, three are predicted to encode non-functional proteins, including a *PPO1-A1* allele with a 54-nucleotide deletion in exon 3 not previously described (Table S3).

Alignment of all 25 *PPO* genes revealed a 38-nucleotide region within the CuB binding domain that shared 100% identity in all seven *PPO1* and *PPO2* target genes (Figure 3). A sgRNA was designed to target the sequence between nucleotides 1,462 and 1,481 on the antisense strand, including a “GGG” Protospacer Adjacent Motif (PAM) (Table S4). Although this PAM was also present in the 18 non-target *PPO* genes, each of these genes contained at least two polymorphisms within the protospacer sequence (Figure 3). No off-target effects are predicted in the protein-coding region of any other gene in the wheat genome and while the promoters of eight genes are potentially targeted, each had at least three mismatches with the protospacer sequence (Table S5). Taken together, this analysis suggests the designed sgRNA should target all seven *PPO1* and *PPO2* genes with minimal off-target activity.

**Figure 3:**
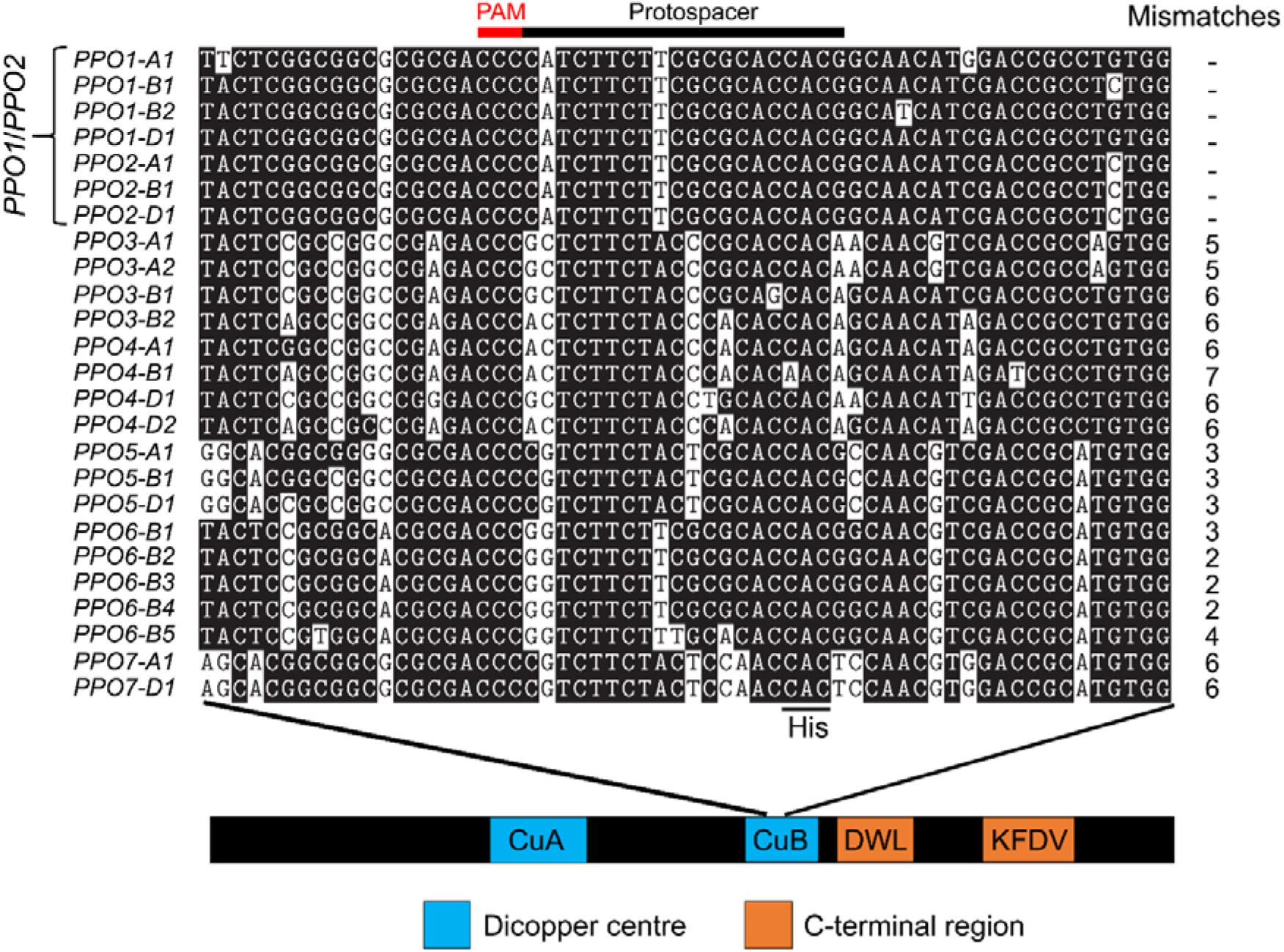
A sgRNA designed to target seven *PPO1* and *PPO2* genes in wheat. The 20-nucleotide protospacer sequence and three nucleotide protospacer adjacent motif (PAM) are indicated. The targeted region encodes the conserved copper binding site II (CuB) domain required for PPO protein function. The codon encoding the third conserved histidine residue (CAC or CAT) is highlighted within the CuB binding domain. Sequence alignments are displayed in 5’-3’ orientation to show the position of the sgRNA, which is designed to the antisense strand. The displayed region is from nucleotides 1,442 to 1,501 based on the distance from the ATG start of the genomic DNA of *PPO1-A1* in ‘Fielder’. The protospacer position is 1,462-1,481 on the reverse strand and is preceded by a ‘GGG’ PAM site. Number of mismatches between the protospacer and each target gene is shown to the right of the alignment.

All seven T_0_ plants regenerated from embryos transformed with the CRISPR/Cas9 construct exhibited induced variation 3-4 nucleotides upstream of the PAM in at least one *PPO* target gene and were selfed to generate T_1_ seed. Of the seven T_1_ lines, five exhibited significant reductions in PPO activity compared to wild-type ‘Fielder’ (*P* < 0.01), ranging from a 45.1% reduction in line 81.5a to an 80.7% reduction in line 81.12a (Figure 4a, Table S6). Grain PPO activity in lines 81.8b and 81.16a were not significantly different from wild-type ‘Fielder’ (*P* >0.05), with the latter line exhibiting higher mean PPO activity than in wild-type ‘Fielder’ (Figure 4a, Table S6).

**Figure 4:**
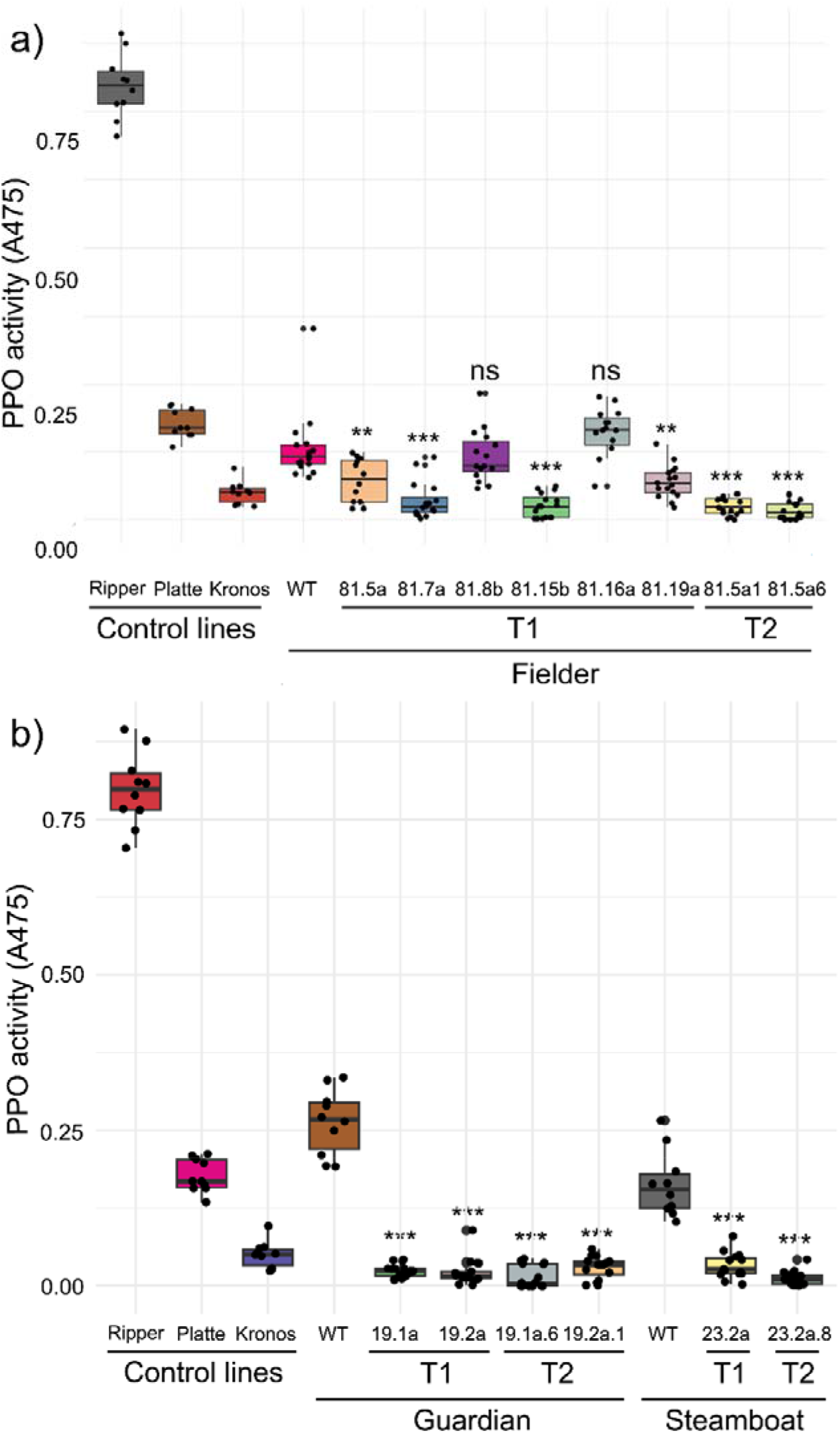
PPO activity in wild-type and edited wheat lines. Mean PPO activity from wild-type, T_1_ and T_2_ lines in a) the spring variety ‘Fielder’ (n = 12 to 18) and b) the winter varieties ‘Steamboat’ and ‘Guardian’. ‘Ripper’ (high-PPO bread wheat), ‘Platte’ (low-PPO bread wheat) and ‘Kronos’ (low-PPO durum wheat) were included as control lines (n = 10). ** = *P* < 0.01. *** = *P* < 0.001. ns = not significant.

The variation in PPO activity both between and within T_1_ lines suggests a complex segregation pattern of edited alleles for each target gene. This was reflected in genotypic data. Two different T_1_ individuals from line 81.5a exhibited examples of different edited alleles of the same target gene (*PPO1-D1*), biallelic edits (*PPO1-A1*), and the absence of edits in some target genes (*PPO1-D2*) (Table S7). Two individuals that exhibited the lowest PPO activity in T_1_ line 81.5a were selfed to generate T_2_ lines. Genotyping of selected T_2_ individuals revealed that they carry a greater number of fixed, non-functional induced alleles than T_1_ plants (Table S7). Mean PPO activity in these T_2_ lines was 80.9% and 86.7% lower than in wild-type ‘Fielder’ (*P* < 0.001) which was greater than the reduction in the corresponding T_1_ lines and with a lower standard deviation (Figure 4a, Table S6).

### Genome editing reduces PPO activity in two elite winter wheat cultivars

The sequence of the sgRNA-PAM is 100% identical in all *PPO1* and *PPO2* genes in fifteen wheat varieties with assembled genomes (Walkowiak et al., 2020), suggesting this construct can be used to edit *PPO1* and *PPO2* genes in diverse wheat germplasm. To test this, the editing construct was transformed into the elite winter wheat cultivars ‘Guardian’ and ‘Steamboat’. No genome assembly is available for these cultivars, so PCR amplification was used to confirm the presence of each *PPO1* and *PPO2* gene. Homoeolog-specific PCR assays for *PPO1-B1* and *PPO1-B2* consistently failed to generate an amplicon in either ‘Guardian’ and ‘Steamboat’, suggesting the absence of these genes in these varieties. *PPO1-B1* and *PPO1-B2* were also absent from eight other common wheat genomes (Table S1) likely because the progenitors of these lines did not carry the duplication event that originated these genes (Figure S1).

Two independent T_0_ ‘Guardian’ plants and one T_0_ ‘Steamboat’ plant exhibited edits in all five target *PPO1* and *PPO2* genes. Derived T_1_ populations from each of these plants all exhibited significant reductions in PPO activity (*P* < 0.001) ranging from an 80.2% reduction in ‘Steamboat’ line 23.2b to a 91.6% reduction in ‘Guardian’ line 19.2a compared to their respective wild-type controls (Figure 4b, Table S6). The reduction in grain PPO activity was even greater in T_2_ lines derived from T_1_ plants exhibiting the lowest PPO activity, including a 96.0% reduction compared to wild-type in ‘Guardian’ line 19.2a.6 and a 92.4% reduction in ‘Steamboat’ line 23.2a.8 (Figure 4b, Table S6). Genotyping of selected individuals confirmed that both T_1_ and T_2_ plants carried induced edits in each target gene (Tables S8 and S9). Mean PPO activity in these T_1_ and T_2_ lines is lower than in the durum wheat ‘Kronos’ (Figure 4b), a genotype that is commonly included as an extremely low-PPO control line and in which only *PPO1-A1* and *PPO2-A1* encode functional PPO enzymes (Table S10). Furthermore, multiple individuals within these T_2_ populations exhibited undetectable PPO activity (Table S6), demonstrating that by editing *PPO1* and *PPO2* genes, it is possible to eliminate grain PPO activity in elite wheat varieties.

## DISCUSSION

### Multi-target genome editing in polyploid wheat

One application of the genome editing tool CRISPR/Cas9 is to simultaneously induce novel genetic variation at multiple loci, including those in the same linkage block. This is especially powerful when targeting multi-gene families such as PPO that are subject to a high rate of gene expansion (Tran et al., 2012). Another recent example is the use of CRISPR/Cas9 to edit multiple ω- and γ-gliadin genes arranged in tandemly duplicated gene clusters (Yu et al., 2023). A growing set of wheat genomes assembled using long-read sequencing data (Walkowiak et al., 2020) facilitates the characterization of this variation and ensures the appropriate design of CRISPR/Cas9 constructs for each target variety. In the current study, the ‘Fielder’ genome (Sato et al., 2021) was used to design a sgRNA targeting a region of the highly conserved CuB binding domain that is 100% identical between all seven target *PPO1* and *PPO2* genes (Figure 3). In addition to facilitating multi-target editing, designing protospacers in a conserved domain increases the likelihood that in-frame deletions or insertions will disrupt gene function. For example, the 15-bp deletion in *PPO1-B2* in ‘Fielder’ (Table S7) eliminates the highly-conserved His and Phe amino acid residues that likely play a critical role in PPO enzyme function (Tran et al., 2012). This contrasts with an earlier CRISPR/Cas9 study to edit *PPO1* genes that used a sgRNA targeting a genomic region between the conserved CuA and CuB binding domains (Zhang et al., 2021). There are polymorphisms between this protospacer sequence and four of the seven *PPO1*/*PPO2* genes from ‘Fielder’, including four mismatches with *PPO2-A1, PPO2-B1,* and *PPO2-D1* (Table S11). These genes are expressed during grain development (Figure 2) and likely contribute to PPO activity in this tissue (Beecher et al., 2012; Liu et al., 2020; Taranto, 2015) suggesting that null alleles in all *PPO1* and *PPO2* genes will be required to maximize reductions in PPO activity by editing.

In the current study, wheat plants with extremely low PPO activity in their grains were developed for all three target varieties that carried disruptive mutations in all *PPO1* and *PPO2* genes, including three individual T_2_ plants with undetectable PPO activity. These observations suggest that despite the presence of *PPO4-D2* transcripts in the developing grain (Figure 2) and genetic studies that identified QTL associated with PPO activity overlapping with *PPO3A-1* and *PPO7D-1* (Liu et al., 2020; Zhai et al., 2020), other *PPO* genes do not make a major contribution to PPO activity in the wheat grain.

### CRISPR/Cas9 design in wheat

It is important to note that while a high rate of editing was achieved using the sgRNA described in the main text, two other sgRNAs targeting a region approximately 150 nucleotides upstream that is also conserved in all seven *PPO1* and *PPO2* genes exhibited zero editing efficiency in 15 T_0_ plants screened for edits (Table S4). It is possible that these CRISPR/Cas9 constructs may have induced transgenerational editing in the T_1_ generation, as observed in previous studies (Wang et al., 2018; Zhang et al., 2021), but this was not evaluated due to the high rate of editing observed with the selected sgRNA. These results strongly suggest major differences in editing efficiencies for sgRNAs targeting DNA sequences in close proximity and that the protospacer sequence composition is critically important for editing efficiency. This is despite the sgRNAs exhibiting comparable “Rule Set 2” (RS2) scores (Table S4), a metric predicting on-target editing efficiency used in CRISPR design tools (Cram et al., 2019; Liu et al., 2017a). This score is derived from models built on empirical editing data from hundreds of constructs used in animal studies (Doench et al., 2016). It is possible that these models do not capture factors that influence editing efficiency in plant species. As the use of CRISPR/Cas9 across different plant species becomes increasingly common, it would be valuable for the plant research community to coordinate editing datasets to develop genus- or species-specific models that can more accurately predict editing efficiency. Highly predictive models would be especially useful when designing editing strategies with a limited number of potential protospacer sequences, such as in multi-target editing or when a highly specific target edit is required.

### Physiological role of PPO in wheat

Despite indirect evidence from some species, the physiological role of PPOs in the plant kingdom remains unclear. Unusually for oxidative enzymes, the size of the *PPO* gene family is highly variable between species and might be driven by clade-specific responses to diverse environmental stresses (Tran et al., 2012). The absence of *PPO* genes from the Arabidopsis genome shows they are not essential and are unlikely to play a role in primary metabolism, but might instead be involved in either environmental responses or secondary metabolism (Tran et al., 2012). In cereals, it has been suggested that PPO activity in grain tissues may contribute to the biochemical resilience to decay in the dormant seed (Fuerst et al., 2014) and in reducing the incidence of black-point, a condition that reduces the quality and aesthetics of wheat products (Liu et al., 2017b). The edited lines exhibiting extremely low PPO activity are ideal near-isogenic materials to test these hypotheses and to characterize the role of *PPO1* and *PPO2* genes in field conditions. In addition, lines carrying different combinations of *PPO1* and *PPO2* null alleles would help determine whether the genes in this family exhibit functional redundancy.

### Applications in breeding

The protospacer sequence used in the current study is 100% conserved in all *PPO1* and *PPO2* genes from 17 wheat genome assemblies screened and conferred significant reductions in PPO activity in all three varieties tested (Figure 4), suggesting this approach can be applied in diverse wheat germplasm. This protospacer sequence is also 100% conserved in orthologous *PPO1* and *PPO2* genes from barley (*Hordeum vulgare*) (Taketa et al., 2010) and rye (*Secale cereale*) (Table S12), so could likely be applied in these species by cloning the sgRNA into an appropriate transformation construct. However, the orthologous *PPO1* and *PPO2* genes from rice (*Oryza sativa*) (Yu et al., 2008), maize (*Zea mays*), and millet (*Sorghum bicolor*) all contained multiple polymorphisms in the protospacer sequence (Table S12). The high conservation of the CuB binding domain in PPO proteins make this region an excellent target for multi-target gene editing in other species, including to generate non-transgenic low-PPO varieties of crops such as mushrooms, potatoes and apples for which RNAi and amiRNA have previously been applied to reduce PPO activity (Chi et al., 2014; González et al., 2019; Murata et al., 2001; Waltz, 2016). This approach may also find application in pea (*Pisum sativum*) and faba bean (*Vicia faba* L.) breeding, where natural null *PPO* alleles conferring a pale hilum colour have been selected in cultivated varieties for their preference by consumers (Balarynová et al., 2022; Jayakodi et al., 2023).

Advances in genotype-independent transformation technologies facilitates genome editing directly in elite wheat cultivars (Debernardi et al., 2020; Wang et al., 2022). Editing *PPO1* and *PPO2* genes in the late stages of variety development would eliminate the need to select for PPO activity in earlier generations, saving breeders time and resources, expand access to high-PPO wheat germplasm, and maximise profits for growers by ensuring high flour yield and quality for all markets. This trait is likely to be especially desirable for applications using whole white wheat flour which retains a higher proportion of aleurone tissue that is removed during white flour refining. It will be necessary to comprehensively phenotype low-PPO edited wheat plants in the field for any undesirable pleiotropic phenotypes, including biotic stress resistance or secondary metabolism. Crosses have been initiated between edited and wild-type plants to generate individuals segregating for the transgene insertion to select edited, non-transgenic lines to phenotype these materials in replicated field trials.

As the application of different CRISPR-derived tools becomes more efficient in different crops and as laws and regulations in some key markets show some signs of loosening (for example, the recent passage of the Genetic Technology (Precision Breeding) Act 2023 through the UK parliament), the question of which alleles to edit becomes more urgent. Evolutionary selection favours mutations in genes with low pleiotropy, that are expressed in a small number of tissues, and which are predicted to be associated with few biological processes (Stern and Orgogozo, 2008). Reported applications of CRISPR/Cas9 in wheat including the *PPO1* and *PPO2* genes described here, as well as *TaASN2* (Raffan et al., 2023) and glutenin genes (Yu et al., 2023) match this profile. An underexplored source of adaptive mutations are gain-of-function alleles that affect transcriptional regulation (Martin and Orgogozo, 2013) as demonstrated previously (Rodríguez-Leal et al., 2017; Song et al., 2022). Identifying beneficial, non-pleiotropic allelic variants that can be directly edited into elite varieties will be essential to fully exploit the power of genome editing for crop improvement.

### Conclusions

Seven *PPO* genes were edited using one sgRNA in hexaploid wheat to generate plants with extremely low grain PPO activity. Directly editing these genes in the late stages of elite variety development may be a complementary approach to accelerate crop improvement, reducing the burden of selecting for multiple loci during early stages of selection. Before these alleles can be deployed in breeding programmes, it will be important to assess the performance of low-PPO edited wheat lines in replicated field experiments to understand the impacts on wheat physiology and performance.

## EXPERIMENTAL PROCEDURES

### Plant materials and growth conditions

The common wheat (*Triticum aestivum* L.) varieties ‘Fielder’, ‘Guardian’, ‘Steamboat’, ‘Ripper’ and ‘Platte’, and the durum wheat (*Triticum turgidum* subsp. *durum* Desf.) variety ‘Kronos’ were used in this study. Seeds of ‘Fielder’ were provided by Dr. David Garvin (USDA-ARS, St. Paul, MN) and seeds of all other varieties were provided by the Colorado State University Wheat Breeding Program. Seeds were germinated in Anchor Paper Co. germination paper for 7 days until emergence, then sown into 1-gallon pots, 2 seedlings per pot, containing water saturated Promix HP Plus Biofungicide and Mycorrhizae potting mix and Osmocote Plus 15-9-12. Two-week-old seedlings of ‘Guardian’ and ‘Steamboat’ were first transferred to plastic bags and vernalized for 6 weeks at 4 °C before being transferred to 1-gallon pots. All plants were grown in greenhouse conditions supplemented by light to maintain a 16 h photoperiod. Temperatures were maintained between 22 □ and 25 □ during the day and between 18 □ and 22 □ during the night. Plants were treated with pesticides as required.

### *PPO* sequence analysis

Genomic DNA sequence of the 20 *PPO* genes previously described (Liu et al., 2020) were extracted from the ‘Chinese Spring’ IWGSC v2.0 reference genome (IWGSC, 2018; Zhu et al., 2021) and used as BLASTn queries to identify *PPO* genes in the assemblies of the spring wheat variety ‘Fielder’ (Sato et al., 2021) and 14 other common wheat varieties (Walkowiak et al., 2020). All PPO genes, including those absent from Chinese Spring but identified in ‘Fielder’, were named following the guidelines endorsed by the Wheat Initiative (Boden et al., 2023). HMMscan was used to confirm the presence of tyrosinase, DWL and KWDV domains in each encoded protein. Microhomology was determined and visualized using the online Triticiae Gene Tribe tool (Chen et al., 2020). The coding sequences of *TaPPO1-A1* and *TaPPO2-A1* were used as queries in BLASTn searches to identify orthologous *PPO* sequences from the genomes of *Hordeum vulgare* (version: MorexV3_pseudomolecules_assembly), *Secale cereale* (version: Rye_Lo7_2018_v1p1p1), *Oryza sativa Japonica* (version: IRGSP-1.0), *Zea mays* (version: Zm-B73-REFERENCE-NAM-5.0), and *Sorghum bicolor* (version: Sorghum_bicolor_NCBIv3).

Expression levels of all *PPO* genes were calculated from mapping a developmental timecourse RNA-seq dataset (Choulet et al., 2014) to the IWGSC v1.2 genome assembly as previously (IWGSC, 2018) (Pearce et al., 2015). Derived transcript per million (TPM) values were displayed as a heatmap using the R package “pheatmap” and scaled for each timepoint using the function “scale(Data_num)”.

### CRISPR/Cas9 plasmid assembly and transformation

The CRISPR design tools CRISPR-P (Liu et al., 2017a) and wheatCRISPR (Cram et al., 2019) were used to support sgRNA design, incorporating “Rule Set 2” scores to estimate editing efficiency (Doench et al., 2016) and scanning the wheat genome to identify potential off-target editing effects. The protospacer was selected based on its high RS2 score, 100% identity to all seven target *PPO1* and *PPO2* genes and low predicted off-target activity in other genes in the wheat genome. A single G nucleotide was added to the start of the 20 nucleotide sgRNA sequence, and the 21-nucleotide sequence (GCGTGGTGCGCGAAGAAGATG) was synthesized as overlapping, complementary oligos with overhanging 5’ and 3’ ends complementary to the insertion site of the target vector. The JD633 vector (Debernardi et al., 2020) was digested with *Aar*Ι and the hybridized oligos were inserted by Golden Gate cloning. This sgRNA was integrated immediately downstream of the U6 promoter. The vector also contains *ZmUbi1*::*SpCas9* and *TaGRF4*:*TaGIF1* coding sequences which confer improved regeneration rates in transformed callus tissue (Debernardi et al., 2020). Ligated vectors were confirmed by Sanger sequencing and transformed into DH5-α *Escherichia coli* cells from which purified plasmid DNA was extracted. After confirming sequence insertion and integrity by Sanger sequencing, plasmid DNA was transformed into *Agrobacterium tumefaciens* strain AGL1 by heat shock and transformed into each wheat genotype using embryo transformation as described previously (Hayta et al., 2021).

### Genotyping

Leaf tissue was harvested from regenerated plants after they had been moved to wheat rooting and growth media (Hayta et al., 2021) and had developed a minimum of 3 leaves at least 4 cm in length. DNA was extracted using the standard CTAB extraction method (Allen et al., 2006) and normalized to 200 ng/µL. Putative transgenic plants were validated by PCR assays to amplify two fragments of the transformed plasmid using the primers listed in Table S13. To characterize induced edits, homoeolog-specific PCR assays were designed to amplify each of the seven target *PPO1* and *PPO2* genes (Table S13). Because of variation in the target sequence, assays for *PPO1-D1* and *PPO2-D1* were customized for different genotypes.

Each PCR consisted of 2.5 µL 10X Standard *Taq* Reaction Buffer (NEB, Ipswich, MA, USA), 0.5 µL 10 mM dNTPs (Invitrogen, Life Technologies, Carlsbad, CA, USA), 0.5 µL 10 µM Forward Primer, 0.5 µL 10 µM Reverse Primer, 5 µL Template DNA (50 ng/µl), 0.125 µL *Taq* DNA Polymerase (NEB, Ipswich, MA, USA) and nuclease-free water to a total reaction volume of 25 µL. For some reactions, HotStarTaq DNA polymerase (Qiagen, Hilden, Germany) was used, using the appropriate buffers and heat activation thermocycler steps recommended by the manufacturer. Reactions were run on a thermocycler using the following conditions: 95 □ 5 min; 40 cycles of 95 □ 30 s, 55-65 □ 30 s, 68 □ 30 s – 2 min; 72 □ 7 min. Annealing temperature and extension time varied by PCR assay and are described with the corresponding primers in Table S13. Selected PCR amplicons were purified using ExoSAP-IT™ PCR Product Cleanup Reagent (Thermo Fisher Scientific, Waltham, MA, USA) and sequenced with Sanger sequencing (Genewiz, Azenta Life Sciences).

### Phenotyping

PPO content was assessed using the L-DOPA method (AACC International Method 22-85.01) using mature harvested wheat grains from greenhouse-grown plants. For each genotype, 5 kernels were placed into a 2 mL microcentrifuge tube before adding 1.5 mL of a solution of 5 mM L-DOPA solution in 50 mM MOPS (pH 6.5). The tubes were sealed, then rotated at 10 rpm for two hours to allow oxygen into the reaction. Absorbance of the resulting solution was measured using 1 mL of sample in a spectrophotometer set to measure at 475 nm using L-DOPA solution as a zero sample. Grains from three other wheat varieties harvested from field experiments in the Colorado State University wheat breeding program were included as controls; the durum wheat (*Triticum turgidum* subsp. *durum*) ‘Kronos’, as an extremely low PPO sample, the winter wheat ‘Ripper’, as a high PPO control, and the winter wheat ‘Platte’, as a low PPO control. Seeds from untransformed wild-type plants of ‘Guardian’, ‘Steamboat’ and ‘Fielder’ were used as comparisons for the corresponding edited lines. The number of replications used for each genotype is listed in Table S6. To determine the significance of differences between lines, pairwise two-tailed Student’s t-tests were applied.

## ACKNOWLEDGEMENTS

We are grateful to Dr. Mervin Poole (Heygates Ltd.) and Dr. Scott Haley (Colorado State University) for helpful discussions about PPO activity in wheat breeding and milling. This project was supported by Competitive Grant 2022-68013-36439 (WheatCAP) from the USDA National Institute of Food and Agriculture and by funding from the Colorado Wheat Administrative Commission and the Colorado Wheat Research Foundation.

## AUTHOR CONTRIBUTIONS

Designed and performed the research: FWM, CK, RAR, KR, AK, JS. Funding acquisition: REM, SP. Project management: REM, SP. Wrote the first draft of the manuscript SP. All authors reviewed and corrected the final text.

## SHORT LEGENDS FOR SUPPORTING INFORMATION

**Table S1:** Presence/absence variation in *PPO1* and *PPO2* genes.

**Table S2:** *PPO* genes in ‘Chinese Spring’ and ‘Fielder’.

**Table S3**: Alleles of *PPO1* and *PPO2* genes in ‘Fielder’.

**Table S4:** Details of three sgRNAs designed to target *PPO1* and *PPO2* genes in wheat.

**Table S5:** Predicted off-target effects of the selected sgRNA used in this study.

**Table S6:** PPO activity in wild-type, T_1_ and T_2_ lines from the varieties ‘Fielder’, ‘Guardian’ and ‘Steamboat’.

**Table S7**: Editing events in seven *PPO1* and *PPO2* genes in selected T_1_ and T_2_ ‘Fielder’ individuals.

**Table S8:** Editing events in seven *PPO1* and *PPO2* genes in selected T_1_ and T_2_ ‘Guardian’ individuals.

**Table S9**: Editing events in seven *PPO1* and *PPO2* genes in selected T_1_ and T_2_ ‘Steamboat’ individuals.

**Table S10:** Alleles of *PPO1* and *PPO2* genes in ‘Kronos’.

**Table S11:** Mismatches between the sgRNA described by Zhang *et al*. 2021 and the seven *PPO1* and *PPO2* genes in the ‘Fielder’ genome.

**Table S12:** Mismatches between the sgRNA used in the current study and orthologous *PPO1* and *PPO2* genes from closely related species.

**Table S13:** Primers used for PCR in the current study.

**Figure S1:** Position of *PPO1* and *PPO2* genes on chromosome 2B in the wheat landrace ‘Chinese Spring’ and the variety ‘CDC Landmark’.

