## Supplemental materials for "Multi-target genome editing reduces polyphenol oxidase activity in wheat (*Triticum aestivum* L.) grains"

This document includes Tables S1 to S13 and Figure S1:

**Table S1**: Presence/absence variation in *PPO1* and *PPO2* genes.

**Table S2:** *PPO* genes in 'Chinese Spring' and 'Fielder'.

**Table S3**: Alleles of *PPO1* and *PPO2* genes in 'Fielder'.

**Table S4:** Details of three sgRNAs designed to target *PPO1* and *PPO2* genes in wheat.

**Table S5:** Predicted off-target effects of the selected sgRNA used in this study.

**Table S6:** PPO activity in wild-type, $T_1$ and $T_2$ lines from the varieties 'Fielder', 'Guardian' and 'Steamboat'.

**Table S7**: Editing events in seven *PPO1* and *PPO2* genes in selected $T_1$ and $T_2$ 'Fielder' individuals.

**Table S8**: Editing events in seven *PPO1* and *PPO2* genes in selected $T_1$ and $T_2$ 'Guardian' individuals.

**Table S9**: Editing events in seven *PPO1* and *PPO2* genes in selected $T_1$ and $T_2$ 'Steamboat' individuals.

**Table S10**: Alleles of *PPO1* and *PPO2* genes in 'Kronos'.

**Table S11:** Mismatches between the sgRNA described by Zhang *et al*. 2021 and the seven *PPO1* and *PPO2* genes in the 'Fielder' genome.

**Table S12:** Mismatches between the sgRNA used in the current study and orthologous *PPO1* and *PPO2* genes from closely related species.

**Table S13:** Primers used for PCR in the current study.

**Figure S1:** Position of *PPO1* and *PPO2* genes on chromosome 2B in the wheat landrace 'Chinese Spring' and the variety 'CDC Landmark'.

**Table S1**: Presence/absence variation in *PPO1* and *PPO2* genes in 17 common wheat varieties.

| Variety | *PPO1-A1* | *PPO2-A1* | *PPO1-B1* | *PPO1-B2* | *PPO2-B1* | *PPO1-D1* | *PPO2-D1* |
|---|---|---|---|---|---|---|---|
| Arina | Present | Present | Present | Present | Present | Present | Present |
| Chinese Spring | Present | Present | Present | Present | Present | Present | Present |
| Claire | Present | Present | Present | Present | Present | Present | Present |
| Fielder | Present | Present | Present | Present | Present | Present | Present |
| Jagger | Present | Present | Present | Present | Present | Present | Present |
| Norin 61 | Present | Present | Present | Present | Present | Present | Present |
| Robigus | Present | Present | Present | Present | Present | Present | Present |
| Lancer | Present | Present | Present | Present | Absent | Present | Present |
| Julius | Present | Present | Present | Present | Absent | Present | Present |
| Cadenza | Present | Present | Absent | Absent | Present | Present | Present |
| Landmark | Present | Present | Absent | Absent | Present | Present | Present |
| Mace | Present | Present | Absent | Absent | Present | Present | Present |
| Paragon | Present | Present | Absent | Absent | Present | Present | Present |
| Stanley | Present | Present | Absent | Absent | Present | Present | Present |
| SY Mattis | Present | Present | Absent | Absent | Present | Present | Present |
| Guardian | Present | Present | Absent | Absent | Present | Present | Present |
| Steamboat | Present | Present | Absent | Absent | Present | Present | Present |

**Table S2:** *PPO* genes in 'Chinese Spring' and 'Fielder'. Gene names are presented according to the guidelines of the Wheat Initiative (Boden et al., 2023) and consistent with previously described allelic variation. Previously proposed gene names are listed (Liu et al., 2020), alongside gene IDs from Chinese Spring (IWGSC v1.2) and Fielder (v1) genome assemblies. Genes that are absent from the 'Chinese Spring' genome assembly are marked with a dash (-). The seven target *PPO1* and *PPO2* genes are separated from non-target *PPO* genes by a horizontal line.

| Gene name | Previous gene name (Liu et al. 2020) | Chinese Spring | Fielder |
|---|---|---|---|
| *PPO1-A1* | *PPO2A-1* | *TraesCS2A02G468200* | *TraesFLD2A01G514000* |
| *PPO1-B1* | *PPO2B-1* | *TraesCS2B02G491000* | *TraesFLD2B01G529900* |
| *PPO1-B2* | *PPO2B-2* | *TraesCS2B02G491100* | *TraesFLD2B01G530000* |
| *PPO1-D1* | *PPO2D-1* | *TraesCS2D02G468200* | *TraesFLD2D01G517900* |
| *PPO2-A1* | *PPO2A-2* | *TraesCS2A02G468500* | *TraesFLD2A01G514300* |
| *PPO2-B1* | *PPO2B-3* | *TraesCS2B02G491400* | *TraesFLD2B01G530300* |
| *PPO2-D1* | *PPO2D-2* | *TraesCS2D02G468600* | *TraesFLD2D01G518200* |
| *PPO3-A1* | *PPO3A-1* | *TraesCS3A02G326600* | *TraesFLD3A01G347800* |
| *PPO3-A2* | - | - | *TraesFLD3A01G347700* |
| *PPO3-B1* | *PPO3B-1* | *TraesCS3B02G355900* | *TraesFLD3B01G404300* |
| *PPO3-B2* | *PPO3B-2* | *TraesCS3B02G568600* | *TraesFLD3B01G637100* |
| *PPO4-A1* | *PPO5A-1* | *TraesCS5A02G549200* | *TraesFLD5A01G586800* |
| *PPO4-B1* | *PPO4B-1* | *TraesCS4B02G385600* | *TraesFLD4B01G423700* |
| *PPO4-D1* | *PPO4D-1* | *TraesCS4D02G358500* | *TraesFLD4D01G404500* |
| *PPO4-D2* | *PPO4D-2* | *TraesCS4D02G359200* | *TraesFLD4D01G405400* |
| *PPO5-A1* | *PPO6A-1* | *TraesCS6A02G041900* | *TraesFLD6A01G054300* |
| *PPO5-B1* | *PPO6B-2* | *TraesCS6B02G056900* | *TraesFLD6B01G067500* |
| *PPO5-D1* | *PPO6D-1* | *TraesCS6D02G048400* | *TraesFLD6D01G053200* |
| *PPO6-B1* | *PPO6B-1* | *TraesCS6B02G016400* | *TraesFLD6B01G018100* |
| *PPO6-B2* | - | - | *TraesFLD6B01G017900* |
| *PPO6-B3* | - | - | *TraesFLD6B01G018000* |
| *PPO6-B4* | - | - | *TraesFLD6B01G018200* |
| *PPO6-B5* | - | - | *TraesFLD6B01G017300* |
| *PPO7-A1* | *PPO7A-1* | *TraesCS7A02G054500* | *TraesFLD7A01G056900* |
| *PPO7-D1* | *PPO7D-1* | *TraesCS7D02G049700* | *TraesFLD7D01G064600* |

**Table S3**: Alleles of *PPO1* and *PPO2* genes in 'Fielder'. Wild-type alleles are predicted to encode full-length, functional PPO proteins with all conserved domains.

| Gene name | Fielder Gene ID | Polymorphism | Effect |
|---|---|---|---|
| *PPO1-A1* | *TraesFLD2A01G514000* | 54-bp deletion in third exon (1662_1713del) | Frame shift, stop codon introduced at amino acid 423. |
| *PPO1-B1* | *TraesFLD2B01G529900* | Wild-type | |
| *PPO1-B2* | *TraesFLD2B01G530000* | 1-bp deletion in third exon (1977delC) | Frame shift, stop codon introduced at amino acid 525. |
| *PPO1-D1* | *TraesFLD2D01G517900* | Wild-type | |
| *PPO2-A1* | *TraesFLD2A01G514300* | Wild-type | |
| *PPO2-B1* | *TraesFLD2B01G530300* | Wild-type | |
| *PPO2-D1* | *TraesFLD2D01G518200* | 1-bp substitution in first exon (20C>A) | Stop codon introduced at amino acid 7. |

**Table S4:** Details of three sgRNAs designed to target *PPO1* and *PPO2* genes in wheat. Quality scores include: RS2 - Rule Set 2 quality score derived from the algorithm described by (Doench et al., 2016). Overall score - weighted average of RS2 and cutting frequency determination from the wheatCRISPR tool (Cram et al., 2019). CRISPRP score - quality score determined by the CRISPR-P 2.0 tool (Liu et al., 2017). The total number of $T_0$ plants represents the number of independently-derived plants confirmed by PCR to carry the transgene. A $T_0$ plant was considered "edited" when induced edits in at least one target *PPO1* or *PPO2* gene were detected by Sanger sequencing. All results described in this manuscript used the first sgRNA in this table.

| Position (IWGSC v1.2) | Strand | Sequence | PAM | Overall score | RS2 | CRISPRp score | $T_0$ plants | |
|---|---|---|---|---|---|---|---|---|
| | | | | | | | Total | Edited |
| chr2A: 712,188,023 | Antisense | CGTGGTGCGCGAAGAAGATG | GGG | 0.58 | 0.63 | 0.44 | 10 | 10 |
| chr2A: 712,188,158 | Antisense | GGTCCGGCTGGTCGCCGGCG | CGG | 0.57 | 0.60 | 0.30 | 4 | 0 |
| chr2A: 712,188,155 | Sense | GCCGGCGACCAGCCGGACCC | GGG | 0.51 | 0.34 | 0.06 | 11 | 0 |

**Table S5:** Predicted off-target effects of the selected sgRNA used in this study (CGTGGTGCGCGAAGAAGATG). Cutting Frequency Determination (CFD) represents the predicted cutting efficiency of an off-target sequence relative to the on-target sequence, ranging from 0 (no predicted activity), to 1.0 (full predicted activity). The first row represents the sgRNA selected sequence targeting all seven *PPO1* and *PPO2* genes (CFD = 1). The gene IDs of other genes targeted in the promoter are listed. Off-target effects were predicted using the wheatCRISPR tool (Cram et al., 2019).

| Protospacer sequence | PAM | CFD | Mismatches | Coding region | Promoter | Other genic region | Intergenic | Targeted gene |
|---|---|---|---|---|---|---|---|---|
| CGTGGTGCGCGAAGAAGATG | GG | 1 | 0 | 7 | 0 | 0 | 0 | *TraesCS2A02G468200*<br>*TraesCS2A02G468500*<br>*TraesCS2B02G491000*<br>*TraesCS2B02G491100*<br>*TraesCS2B02G491400*<br>*TraesCS2D02G468200*<br>*TraesCS2D02G468600* |
| TGTGGCACGCGAAGAAGATG | GG | 0.91 | 3 | 0 | 1 | 0 | 0 | *TraesCS7B02G293100* |
| TGTGGCACGTGAAGAAGATG | GG | 0.86 | 4 | 0 | 1 | 0 | 0 | *TraesCS4B02G141200* |
| CATGGCACGTGAAGAAGATG | GG | 0.72 | 4 | 0 | 2 | 2 | 0 | *TraesCS4A02G553800LC*<br>*TraesCS7B02G226300* |
| CGTGAAGCACGAAGAAGATG | GG | 0.48 | 3 | 0 | 0 | 0 | 1 | |
| CGGGGCACGCGAAGAAGATG | GG | 0.46 | 3 | 0 | 0 | 0 | 1 | |
| CGTCGTGTGCAAAGAAAATG | GG | 0.43 | 4 | 0 | 1 | 0 | 0 | *TraesCS3D02G054900* |
| TGAGGTCTGCGAAGAAGATG | GG | 0.43 | 4 | 0 | 1 | 0 | 0 | *TraesCS7A02G282200* |
| CCTAGTCCGCGAAGAAAATG | GG | 0.4 | 4 | 0 | 1 | 0 | 0 | *TraesCS1A02G195500* |
| GCTGGAGCGCGAAGAAGATT | GG | 0.38 | 4 | 0 | 1 | 0 | 0 | *TraesCS6B02G572000LC* |

**Table S6:** PPO activity in wild-type, $T_1$ and $T_2$ lines from the varieties 'Fielder', 'Guardian' and 'Steamboat'. Mean PPO activity, the number of biological replicates (n), standard deviation and two-tailed Student's T-test result showing the difference with each corresponding wild-type line are shown.

| Genotype | Line | Generation | Mean PPO activity (A475) | % of wild-type | n | Standard deviation | *P* value |
|---|---|---|---|---|---|---|---|
| Ripper | Wild-type | Wild-type | 0.797 | - | 10 | 0.019 | |
| Platte | Wild-type | Wild-type | 0.177 | - | 10 | 0.008 | |
| Kronos | Wild-type | Wild-type | 0.050 | - | 10 | 0.007 | |
| Fielder | Wild-type | Wild-type | 0.132 | - | 16 | 0.064 | |
| | 81.5a | $T_1$ | 0.073 | 54.9 | 12 | 0.040 | 0.0058 |
| | 81.7a | $T_1$ | 0.035 | 26.7 | 16 | 0.033 | 2.14 $e^{-5}$ |
| | 81.8b | $T_1$ | 0.117 | 88.5 | 15 | 0.048 | 0.46 |
| | 81.12a | $T_1$ | 0.026 | 19.3 | 16 | 0.022 | 5.86 $e^{-6}$ |
| | 81.15b | $T_1$ | 0.044 | 33.3 | 16 | 0.027 | 6.12 $e^{-6}$ |
| | 81.16a | $T_1$ | 0.161 | 121.7 | 15 | 0.044 | 0.16 |
| | 81.19a | $T_1$ | 0.070 | 52.8 | 16 | 0.030 | 0.0021 |
| | 81.5a.1 | $T_2$ | 0.025 | 19.1 | 17 | 0.016 | 6.42 $e^{-6}$ |
| | 81.5a.6 | $T_2$ | 0.018 | 13.8 | 17 | 0.017 | 2.91 $e^{-6}$ |
| Guardian | Wild-type | Wild-type | 0.263 | - | 10 | 0.052 | |
| | 19.1a | $T_1$ | 0.023 | 8.7 | 17 | 0.011 | 1.00 $e^{-7}$ |
| | 19.2a | $T_1$ | 0.022 | 8.4 | 18 | 0.020 | 3.91 $e^{-8}$ |
| | 19.1a.6 | $T_2$ | 0.014 | 5.1 | 16 | 0.018 | 3.90 $e^{-8}$ |
| | 19.2a.1 | $T_2$ | 0.029 | 11.1 | 16 | 0.018 | 6.20 $e^{-8}$ |
| Steamboat | Wild-type | Wild-type | 0.163 | - | 10 | 0.053 | |
| | 23.2a | $T_1$ | 0.032 | 19.6 | 15 | 0.020 | 1.34 $e^{-5}$ |
| | 23.2a.8 | $T_2$ | 0.012 | 7.6 | 16 | 0.011 | 1.73 $e^{-5}$ |

**Table S7**: Editing events in seven *PPO1* and *PPO2* genes in selected $T_1$ and $T_2$ 'Fielder' individuals. Where mutations are heterozygous, both allele types are described. WT = wild-type allele. * = natural allele is expected to encode a non-functional protein in Fielder (Table S3).

| **$T_1$** | **81.5a.1** | **81.5a.6** |
|---|---|---|
| *PPO1-A1** | [1464_1465delTC]/[1465delC] | [1464_1465delTC] |
| *PPO2-A1* | [1274_1275insC]/[1273_1274delTC] | [1274_1275insC] |
| *PPO1-B1* | [1512_1518delATCTTC] | [1512_1518delATCTTC] |
| *PPO1-B2** | [1369_1383delTTCTTCGCGCACCAC] | [1369_1383delTTCTTCGCGCACCAC] |
| *PPO2-B1* | [1295_1296insT] | [1295_1296insT] |
| *PPO1-D1* | [1273_1274delTC] | [1270subA;1274delC] |
| *PPO2-D1** | No edits | [1266_1267insG;1266_1279delTTCTTCGCGCACC] |
| **$T_2$** | **81.5a.1.6; 81.5a.1.5; 81.5a.1.8.** | **81.5a.6.1; 81.5a.6.4; 81.5a.6.7; 81.5a.6.13.** |
| *PPO1-A1** | [1465delC] | [1464_1465delTC] |
| *PPO2-A1* | [1273_1274delTC] | [1274_1275insC] |
| *PPO1-B1* | [1512_1518delATCTTC] | [1512_1518delATCTTC] |
| *PPO1-B2** | [1369_1383delTTCTTCGCGCACCAC] | [1369_1383delTTCTTCGCGCACCAC] |
| *PPO2-B1* | [1295_1296insT] | [1295_1296insT] |
| *PPO1-D1* | [1273_1274delTC] | [1270subA;1274delC] |
| *PPO2-D1** | No edits | [1266_1267insG;1266_1279delTTCTTCGCGCACC] |

$T_2$ line 81.5a.1.11 has the same genotype, except biallelic alleles in *PPO1-A1* and *PPO1-A2*.

**Table S8**: Editing events in five *PPO1* and *PPO2* genes in selected $T_1$ and $T_2$ 'Guardian' individuals. Where mutations are heterozygous, both allele types are described, separated by "/".

| **$T_1$** | **19.1a.6** | **19.2a.1** |
|---|---|---|
| *PPO2A-1* | [1299_1300insA]/[1296_1304delCCATCTTCT] | [1299_1300insA]/[1299_1300insT] |
| *PPO2A-2* | [1230delC]/[1232subT] | [1232_1235delTCTT]/[1230_1234delCTTCT] |
| *PPO2B-3* | [1367_1382delTCTTCTTCGCGCACCA] | [1367_1368insT] |
| *PPO2D-1* | [1341_1342delTC] | [1341_1343delTCT]/[1341_1342insC] |
| *PPO2D-2* | [1263_1270delCATCTTCT] | [1266_1271delTCTTCT]/<br>[1235_1271delCGCGCGACCCCATCTTCTT] |
| **$T_2$** | **19.1a.6.5** | **19.2a.1.3** |
| *PPO2A-1* | [1299_1300insA]/[1296_1304delCCATCTTCT] | [1299_1300insA] |
| *PPO2A-2* | [1230delC]/[1232subT] | [1230_1234delCTTCT] |
| *PPO2B-3* | [1367_1382delTCTTCTTCGCGCACCA] | [1367_1368insT] |
| *PPO2D-1* | [1341_1342delTC] | [1341_1342insC] |
| *PPO2D-2* | [1263_1270delCATCTTCT] | [1266_1271delTCTTCT] |

**Table S9**: Editing events in five *PPO1* and *PPO2* genes in selected $T_1$ and $T_2$ 'Steamboat' individuals. Where mutations are heterozygous, both allele types are described, separated by "/".

| **$T_1$** | **23.2a.8** |
|---|---|
| *PPO2A-1* | [1273_1299delCTTCTTCTCGGCGGCGCGCGACCCCAT] |
| *PPO2A-2* | [1229_1230insA] |
| *PPO2B-3* | [1368delC]/[1368_1369insC] |
| *PPO2D-1* | [1342delC] |
| *PPO2D-2* | [1267_1269delTTC] |
| **$T_2$** | **23.2a.8.15** |
| *PPO2A-1* | [1273_1299delTCTTCTCGGCGGCGCGCGACCCCATCT] |
| *PPO2A-2* | [1229_1230insA] |
| *PPO2B-3* | [1368delC] |
| *PPO2D-1* | [1342delC] |
| *PPO2D-2* | [1267_1269delTTC] |

**Table S10**: Alleles of *PPO1* and *PPO2* genes in 'Kronos'. *PPO* gene sequences were used as queries in BLAST searches against the Kronos Elv1.1 genome assembly.

| Gene name | Fielder Gene ID | Polymorphism | Effect |
|---|---|---|---|
| *PPO1-A1* | *TraesFLD2A01G514000* | Wild-type | |
| *PPO2-A1* | *TraesFLD2A01G514300* | Wild-type | |
| *PPO1-B1* | *TraesFLD2B01G529900* | Deleted | |
| *PPO1-B2* | *TraesFLD2B01G530000* | Deleted | |
| *PPO2-B1* | *TraesFLD2B01G530300* | 1 nucleotide substitution in first exon, (20C>A) | Stop codon introduced at amino acid 7. |

**Table S11:** Mismatches between the sgRNA described by Zhang *et al*. (2021) and the seven *PPO1* and *PPO2* genes in the 'Fielder' genome. This sgRNA matches 100% only with *PPO1-A1*, *PPO1-B2* and *PPO1-D1*. The protospacer sequence (5'-GAAGAAGACGCTGCTGTTCC-3') designed to the forward strand is between positions 1,291 and 1,310 bp downstream of the initiating ATG codon of *PPO1-A1* based on the 'Fielder' gene model. Mismatches are highlighted in bold. PAM sequence is in red.

| Gene name | Fielder gene ID | sgRNA sequence | Number of mismatches |
|---|---|---|---|
| *PPO1-A1* | *TraesFLD2A01G514000* | GAAGAAGACGCTGCTGTTCCTGG | 0 |
| *PPO2-A1* | *TraesFLD2A01G514300* | **C**AAGAAGAC**T**C**G**GCTGTTC**A**TGG | 4 |
| *PPO1-B1* | *TraesFLD2B01G529900* | GAAGAAGACG**T**TGCTGTTCCTGG | 1 |
| *PPO1-B2* | *TraesFLD2B01G530000* | GAAGAAGACGCTGCTGTTCCTGG | 0 |
| *PPO2-B1* | *TraesFLD2B01G530300* | **C**AAGAAGAC**T**C**G**GCTGTTC**A**TGG | 4 |
| *PPO1-D1* | *TraesFLD2D01G517900* | GAAGAAGACGCTGCTGTTCCTGG | 0 |
| *PPO2-D1* | *TraesFLD2D01G518200* | **C**AAGAAGAC**T**C**C**GCTGTTC**A**TGG | 4 |

**Table S12:** Mismatches between the sgRNA used in the current study and orthologous *PPO1* and *PPO2* genes from closely related species. Mismatches are highlighted in bold. PAM sequence is in red. * The mismatch in *ScPPO1* alters the PAM sequence from CCC to CCT, which is still permissible for editing.

| Species | Gene name | Gene ID | sgRNA sequence | Number of mismatches |
|---|---|---|---|---|
| *Triticum aestivum* | *TaPPO1-A1* | *TraesFLD2A01G514000* | CCCCATCTTCTTCGCGCACCACG | 0 |
| *Hordeum vulgare* | *HvPPO1* | *HORVU.MOREX.r3.2HG0194690* | CCCCATCTTCTTCGCGCACCACG | 0 |
| *Hordeum vulgare* | *HvPPO2* | *HORVU.MOREX.r3.2HG0194630* | CCCCATCTTCTTCGCGCACCACG | 0 |
| *Secale cereale* | *ScPPO1* | *SECCE2Rv1G0121020* | CC**T**CATCTTCTTCGCGCACCACG | 1* |
| *Secale cereale* | *ScPPO2* | *SECCE2Rv1G0121070* | CCCCATCTTCTTCGCGCACCACG | 0 |
| *Zea mays* | *ZmPPO1* | *Zm00001eb431220* | CCC**GG**T**G**TTCTTCGCGCACCACG | 3 |
| *Zea mays* | *ZmPPO2* | *Zm00001eb428900* | CCC**GC**TCTTCTTCGCGCACCACG | 2 |
| *Zea mays* | *ZmPPO3* | *Zm00001eb171410* | CCC**GC**TCTTCT**A**C**T**CGCACCACG | 4 |
| *Zea mays* | *ZmPPO4* | *Zm00001eb388240* | CCCCATCTTCTTC**C**CGCACCAC**A** | 2 |
| *Oryza sativa* | *OsPPO1* | *Os04g0624500* | CCC**GG**T**G**TTCTTCGCGCACCACG | 3 |
| *Oryza sativa* | *OsPPO2* | *Os04g0624450* | CCC**GG**T**G**TTCTTCGCGCACCACG | 3 |
| *Sorghum bicolor* | *SbPPO1* | *SORBI_3007G068500* | CCC**GC**TCTTCT**A**C**T**CGCACCACG | 4 |
| *Sorghum bicolor* | *SbPPO2* | *SORBI_3006G181300* | **T**CC**A**CTCTTCTTCGCGCACCACG | 3 |
| *Sorghum bicolor* | *SbPPO3* | *SORBI_3006G181400* | **T**CC**GC**TCTTCTTCGCGCACCACG | 3 |
| *Sorghum bicolor* | *SbPPO4* | *SORBI_3010G192700* | CCCCATCTTCT**A**C**C**CGCACCACG | 2 |

**Table S13:** Primers used for PCR in the current study. To amplify the *PPO1-D1* and *PPO2-D1* genes, different PCR assays were used in different genotypes due to sequence variation.

| Target | Primer Name | Primer Sequence (5'-3') | Annealing temp (°C) | Annealing time (seconds) | Amplicon size (bp) |
|---|---|---|---|---|---|
| *PPO1-A1* | PPO-A1_F5 | AAAATATTGTTAACATAACCACAGAGTT | 57 | 60 | 972 |
| | PPO-A1_R5 | CCAGAGCGGAGCACATTAC | | | |
| *PPO2-A1* | PPO-A2_F2 | GTAACGTACACCACATAATAAATACGC | 57 | 60 | 1,200 |
| | PPO-A2_R2 | GCCTCCTCCTTCTCCTTGTC | | | |
| *PPO1-B1* | PPO-B1_F3 | GACGCCAATATCCCAAGAGA | 61 | 30 | 933 |
| | PPO-B1_R3 | ACCCTGGGCCTCGTCACA | | | |
| *PPO1-B2* | PPO-B2c_F | GACGCGGTACCAAAATCTCTC | 59 | 30 | 493 |
| | PPO-B2_R | TCCTGCGGGATCTCTTGC | | | |
| *PPO2-B1* | PPO-B1_F3 | GACGCCAATATCCCAAGAGA | 60 | 30 | 725 |
| | PPO-B3_R2 | GGCCTCGTCACCGTCAT | | | |
| *PPO1-D1* (Steamboat) | PPO-D1a_F | CCTCAAGAATCCCTAAACAAAATGAG | 59 | 30 | 1,008 |
| | PPO-D1a_R | GACGCGAGAGCGGACCACATTAG | | | |
| *PPO1-D1* (Guardian, Fielder) | PPO-D1bc_F2 | CGGTCATCTACGCCAACAGA | 61 | 30 | 627 |
| | PPO-D1bc_R2 | CCTCGTCGTAGAAGAGGAAGC | | | |
| *PPO2-D1* (Steamboat) | PPO-D2a_F | TGTCGGGTGCCAAGAAGACTA | 65 | 30 | 275 |
| | PPO-D2_R | CCAGTCGGCGTCGGC | | | |
| *PPO2-D1* (Guardian, Fielder) | PPO-D2b_F | TGTCGAGTGCCAAGAAGACTCC | 65 | 30 | 275 |
| | PPO-D2_R | CCAGTCGGCGTCGGC | | | |
| JD633 plasmid backbone | TaU6-promoter_F | TAGGAGGGAATCGAACTAGG | 57 | 60 | 400 |
| | gRNA-scaffold_R | CTTTTTCAAGTTGATAACGG | | | |
| Hygromycin resistance gene | HYG_F | CTATTTCTTTGCCCTCGGACGAGTGC | 55 | 60 | 1,026 |
| | HYG_R | ATGAAAAAGCCTGAACTCACCGCGAC | | | |
| *TaCas9* | TaCDP-1_F | GGCAGCCCGGAGGACAACGAGC | 60 | 60 | 350 |
| | TaCDP-1_R | GAGAGGTCGATCCTCGTCTCGTA | | | |

**Figure S1:** Position of *PPO1* and *PPO2* genes on chromosome 2B in the wheat landrace 'Chinese Spring' and the variety 'CDC Landmark'. Gene positions are drawn to scale and homologous genes linked by lines determined using the Triticeae Gene Tribe microhomology tool (Chen et al., 2020).

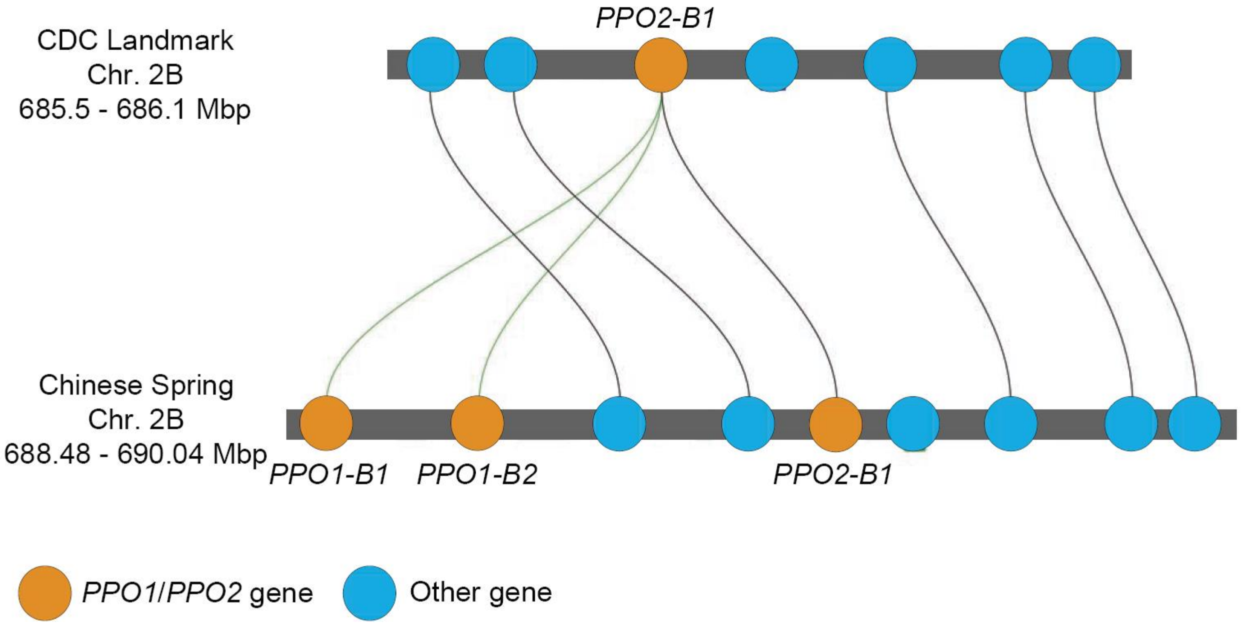
